# Cell-free chromatin particles from dying cells promote the induction of an immune response in human lymphocytes

**DOI:** 10.1101/2023.08.07.551233

**Authors:** Snehal Shabrish, Gorantla V. Raghuram, Nimisha Raphael, Ranjani Narayanan, Relestina Lopes, Dipali Kondhalkar, Naveen Kumar Khare, Indraneel Mittra

## Abstract

It is long established that cell death and immune response are closely related, although the nature of this relationship has remained unclear. We earlier reported that cell-free chromatin particles (cfChPs) released from the billions of cells that die in the body every day to enter into the blood circulation, or those that are released locally from dying cells, can readily enter into healthy cells to induce DNA damage and activate inflammatory cytokines. In this study we investigated whether cfChPs from dying cells might be the missing link between cell death and immune response. We treated human lymphocytes with cfChPs isolated from sera of healthy individuals or with cfChPs released from hypoxia-induced dying lymphocytes. We observed that cfChPs from both sources readily entered into lymphocytes to accumulate in their nuclei within 2 h. This was associated with marked activation of CD69 and release of inflammatory cytokines, as well as of p-STING and cGAS expression. The addition of the STING protein inhibitor H151 to the cfChPs treated cells abolished the release of inflammatory cytokines. These findings lead us to suggest that cfChPs from dying cells are the critical triggers of immune response which act via cGAS-STING pathway.

## Introduction

The mechanism that triggers immune response is a central but an unresolved question in immunology, although there is a well-established association between immune response and cell death (Ashida et al., 2011). Cell death results in the release of cellular components which are thought to act as damage-associated molecular patterns (DAMPs) to activate immune response (Akira et al., 2006; Chaplin, 2010; Land, 2015; Rock et al., 2011). Optimal levels of DAMPs in healthy individuals are required to maintain immune homeostasis; however, in pathological conditions such as sepsis, cancer, neurodegenerative disorders, and auto-inflammatory as well as autoimmune diseases, excessive cell death leads to an uncontrolled release of DAMPs that can provoke hyper-inflammatory responses (Denning et al., 2019; Land, 2015; Rock et al., 2011). However, despite intensive research, the precise component of DAMPs that might be critical for mounting immune response continues to remain obscure, hindering the development of effective therapies for life-threatening disorders (Fleischmann et al., 2016; Schaefer, 2014; Ye et al., 2020).

We have previously reported that cell-free chromatin particles (cfChPs) that are released from dying cells are readily internalized by healthy cells wherein they induce DNA damage and inflammation (Indraneel Mittra et al., 2017; Mittra et al., 2015). We have also shown that cfChPs play an important role in disorders associated with hyper-inflammation, such as toxicity from chemo- and radiotherapy, sepsis, ageing, and cancer (Agarwal et al., 2022; Kirolikar et al., 2018; I. Mittra et al., 2017; Mittra Id et al., 2020; Ostwal et al., 2022; Pal et al., 2022; Pilankar et al., 2022). Since inflammation is a critical component of immune response, these observations led us to hypothesize that cfChPs released from dying cells could be the elusive agents that could link cell death with an inflammatory immune response (Shabrish and Mittra, 2021). In pursuit of this hypothesis, we utilized two experimental models: 1) a culture system wherein peripheral blood mononuclear cells (PBMCs) were exposed to conditioned medium (CM) containing cfChPs released from hypoxia-induced dying PBMCs, and 2) direct treatment of PBMCs with cfChPs isolated from healthy human sera. The immunological effects of cfChPs were investigated by evaluating three integral aspects of an inflammatory immune response viz. 1) immune cell activation characterized by CD69 expression on T cells and NK cells, 2) cellular stress response, and 3) activation of inflammatory cytokines.

## Results

### cfChPs released from dying cells are readily internalized by PBMCs

As a first step, we revisited our previous finding that cfChPs released from dying cells are readily internalized by mouse fibroblast cells (Indraneel Mittra et al., 2017) and confirmed that cfChPs released from dying PBMCs are also internalized by healthy PBMCs. To demonstrate this, PBMCs were dually fluorescently labelled in their DNA and histones using bromodeoxyuridine (BrdU) and histone-2B-GFP, respectively (see Materials and Methods), and induced to undergo cell death by hypoxia. The CM of the dying PBMCs, when added to fresh PBMCs, showed copious accumulation of fluorescently dually labelled cfChPs in their nuclei at 4 h **(Suppl. Fig. 1).**

### Immunogenic effects of CM containing cfChPs

We next investigated the possible immunogenic effects of cfChPs that are contained in CM by evaluating CD69 expression, activation of cellular stress markers, and cytokine release by treating fresh PBMCs with hypoxic CM. To determine the optimum volume of CM required, a dose-response analysis was initially performed, which showed that maximum CD69 expression on T- and NK-cells was achieved with 200 μL CM **(Fig. 1A).** All further experiments were therefore performed using 200 µl CM. The treatment of PBMCs with 200 µL CM was also associated with marked activation of the inflammatory cytokines IL-1β, IL-6, TNF-α, IFN-γ, and CXCL9 **(Fig. 1B).** Since stress-related proteins interact with and regulate signaling intermediates involved in an immune response, we next investigated whether cfChPs from dying cells would activate the cellular stress-related markers c-Jun, c-Fos, JunB, FosB, EGR-1, and NF-КB. Activation of stress markers was evaluated utilizing qRT-PCR in a time course experiment, which showed upregulation of all six stress markers albeit at different time points **(Suppl. Fig. 2)**. Of these, we could confirm the expression of FosB at a protein level by flow cytometry when examined at the maximum time-point of its mRNA expression **(Fig. 1C)**.

**Fig. 1.**
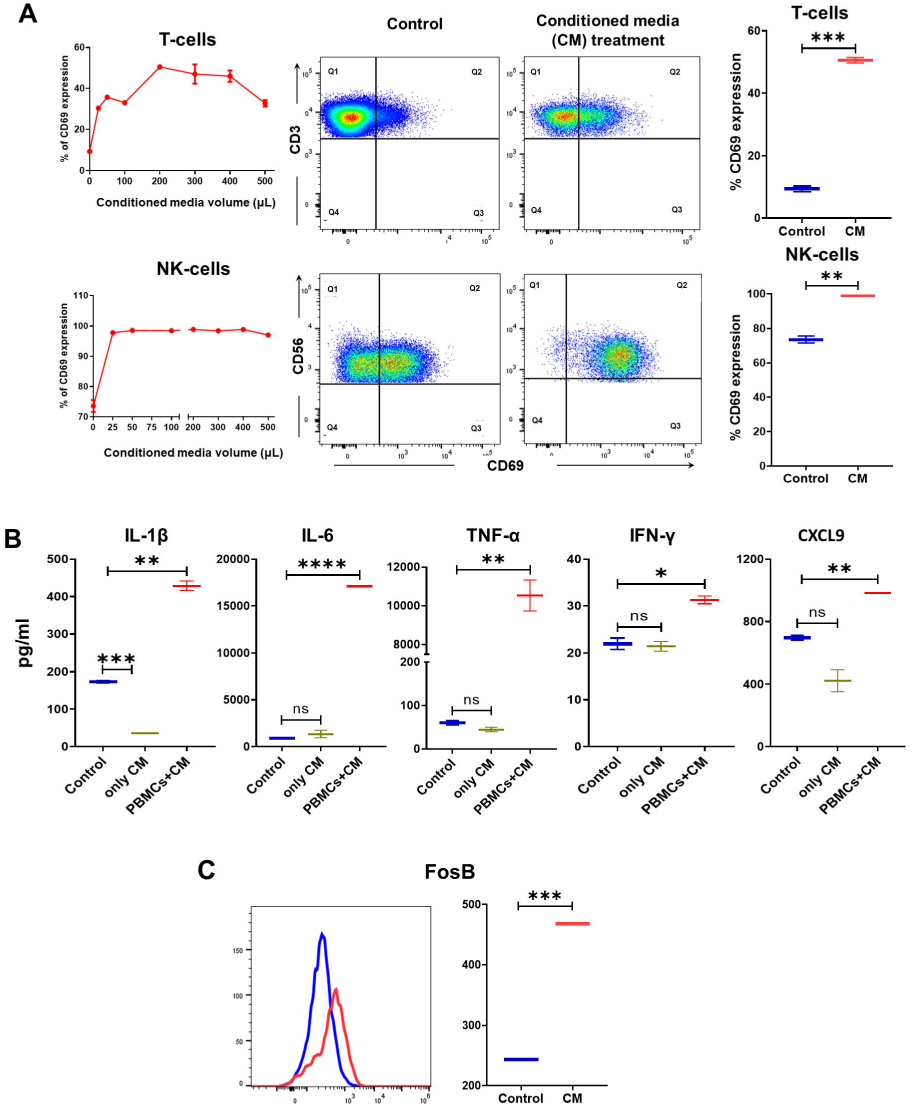
cfChPs released from dying PBMCs into conditioned medium (CM) induce immune response. **(A)** Representative flow cytometry plots of CM-treated PBMCs. Line graphs represent the expression of CD69 in CM-treated PBMCs through dose-response analyses (left hand panel). Representative flow cytometry plots (middle panel) and box-plot (right hand panel) represent the relative upregulation of CD69 expression on T-cells and NK-cells using 200 μL CM. **(B)** Box-plot represent the relative production of inflammatory cytokines by PBMCs in response to CM (200 µl) as estimated by flow cytometry and ELISA. It is to be noted that CM itself does not contain cytokines. **(C)** Upregulation of stress related marker expression in PBMCs treated with CM. Samples were analyzed by flow cytometry and respective protein expression was analyzed on untreated (blue line) and CM treated (red line) PBMCs. Box-plot represents quantitative analysis (MFI) of stress related markers by PBMCs in response to CM. Box-plot represent median with minimum and maximum data values. Statistical analyses were performed using two-tailed unpaired student’s *t*-tests in GraphPad Prism 8. **_*_** p<0.05, **_**_** p<0.01, _***_ p<0.005.

### Serum derived cfChPs are readily internalized by PBMCs

We previously reported that cfChPs isolated from human sera are readily internalized by mouse fibroblast cells (Mittra et al., 2015). Microarray analysis had suggested that cellular uptake of cfChPs likely occurs via the phagocytosis pathway (Indraneel Mittra et al., 2017). Other publications have also shown that nucleosome uptake happens via endocytosis (Wang et al., 2021). Thus, we speculated, whether serum derived cfChPs, like those contained in hypoxic CM, might also activate inflammatory cytokines? As a first step, we reconfirmed that serum derived cfChPs are internalized by PBMCs. We labelled the serum derived cfChPs in their DNA and histones with Pt Bright 550 (red) and ATTO-488 (green; see Materials and Methods), respectively. When the dually labelled cfChPs (10 ng) were added to fresh PBMCs, co-localizing red and green fluorescence signals representing cfChPs were clearly detected in nuclei of fifty percent of the treated PBMCs by 2 h **(Suppl. Fig. 3)**.

### Immunogenic effects of serum derived cfChPs

To investigate whether serum derived cfChPs, like those released from dying cells, would trigger immune response, we treated PBMCs with serum derived cfChPs (10 ng) and evaluated their immunogenic potential. The treatment of PBMCs with cfChPs triggered a marked increase in the expression of CD69 on T-cells and NK-cells at 24 h **(Fig. 2A).** A pronounced increase in levels of the pro-inflammatory cytokines IL-1β, IL-6, TNF-α, IFN-γ, CXCL9, and IFN-β, as well as the two anti-inflammatory cytokines IL-10 and TGF-β1 **(Fig. 2B)**, was also detected. cfChPs also activated mRNA expression NF-КB, c-Jun, c-Fos, Jun-B, FosB, and EGR-1 on human PBMCs, but at different time points **(Suppl. Fig. 4).** Of these, upregulation of four markers namely, c-Jun, c-Fos, Jun-B and FosB were also found to be upregulated at protein level when analyzed by flow cytometry at the peak time point of mRNA expression **(Fig. 2C)**.

**Fig. 2.**
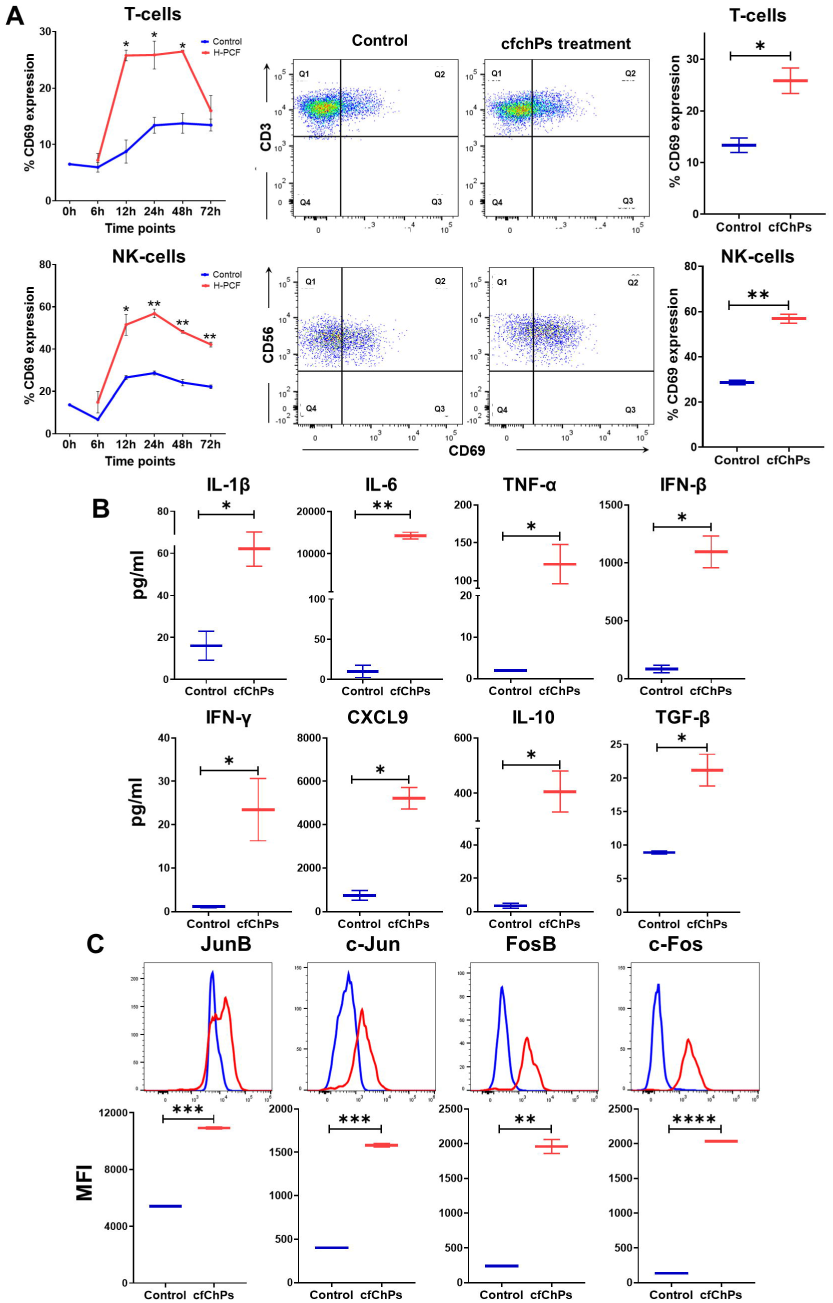
Serum derived cfChPs activate immune response. **(A)** Representative flow cytometry plots of cfChPs (10 ng)-treated PBMCs. Line graphs represent the expression of CD69 in a time course experiment (blue line represents untreated cells; red line represents cfChP-treated cells) (left hand panel). Representative flow cytometry plots (middle panel) and box-plot (right hand panel) represent the relative upregulation of CD69 expression on T-cells and NK-cells in response to cfChPs at the peak time-point (24 h). **(B)** Box-plots representing the production of cytokines by PBMCs in response to cfChPs (10 ng). **(C)** Upregulation of stress related marker expression in PBMCs treated with cfChPs. Samples were analyzed by flow cytometry and respective protein expression was analyzed on untreated (blue line) and cfChPs treated (red line) PBMCs (upper panel). Box-plots in the lower panel represent quantitative analysis (MFI) of stress related markers by PBMCs in response to cfChPs (10 ng). Box-plot represent median with minimum and maximum data values and the data were analyzed using two-tailed unpaired student’s *t*-tests in GraphPad Prism 8. **_*_** p<0.05, **_**_** p<0.01, **_***_** p<0.005, **_****_** p<0.0001.

### Serum derived cfChPs act via the cGAS-STING pathway

Since the cGAS-STING pathway is activated when DNA is atypically present in the cytoplasm (Motwani et al., 2019), we hypothesized that cGAS-STING might be activated when cfChPs appear in the cytoplasm. We investigated this indirectly by treating PBMCs with cfChPs in the presence of the STING protein inhibitor H151 and demonstrate a significantly reduced production of inflammatory cytokines as estimated by flow cytometry and ELISA **(Fig. 3A)**. qRT-PCR performed 12h after treatment of PBMCs with serum derived cfChPs (10 ng) provided direct evidence of the involvement of cGAS-STING pathway **(Fig. 3B)**. Further direct evidence came from our demonstration that cfChPs treatment significantly upregulated phosphorylated STING protein (p-STING) expression in lymphocytes at 24 h as determined by flow cytometry **(Fig. 3C)**. We further show using immunofluorescence that uptake of cfChPs (labelled in their histone with ATTO-488) markedly increased cGAS expression in PBMCs by 4h **(Fig. 3D)**. cfChPs were labelled only in their histones with ATTO-488 (green) to prevent spectral overlap with cGAS staining. Interestingly, in response to the internalized cfChPs, cGAS was found to shuttle between the cytoplasm and nucleus. At 2h of cfChPs treatment, cGAS translocated from the nucleus to the cytoplasm, while at 4 h it had translocated back to the nucleus, indicating that cGAS is directly involved in cfChPs induced immune response. These findings suggest that cfChPs activate immune response via the cGAS-STING pathway.

**Fig. 3.**
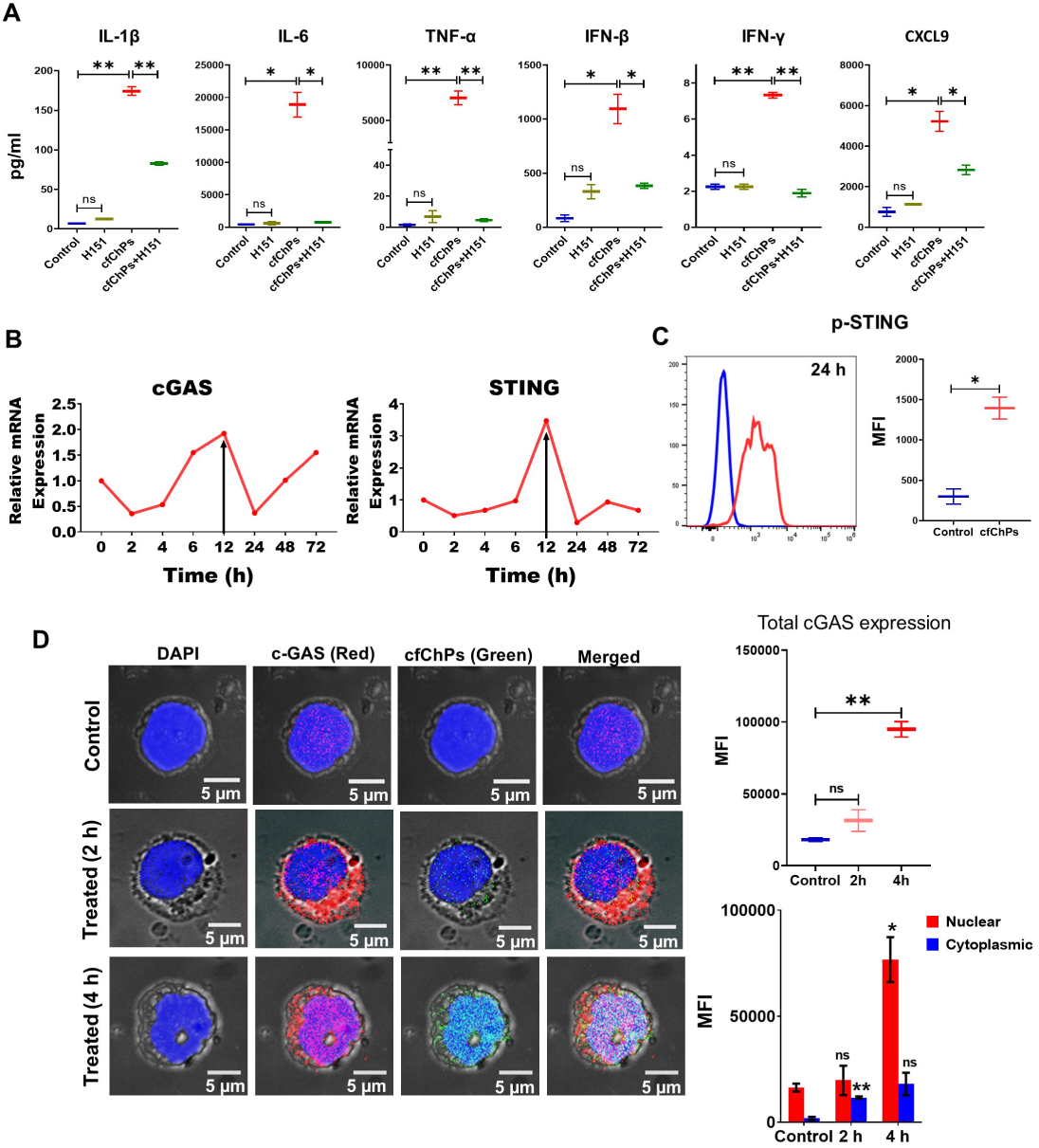
Serum derived cfChPs activate immunogenic effects *via* the cGAS-STING pathway. **(A)** Box-plots represent data of inflammatory cytokines secreted by PBMCs in response to cfChPs (10 ng) and their inhibition by prior treatment with the STING protein inhibitor H151 (50 ng/ml). It is to be noted that treatment with H151 itself had no effects. **(B)** Upregulation of mRNA expression of cGAS and STING in human PBMCs following treatment with serum derived cfChPs (10 ng) detected by qRT-PCR and analyzed using a comparative C_T_ method. Line graphs represent the results of a time course experiment calculated as expression fold [2^(-ΔΔC_T_)]. **(C)** Upregulation of p-STING expression in PBMCs following treatment with serum derived cfChPs (10 ng) detected by flow cytometry (blue line represents untreated cells; red line represents cfChPs-treated cells). Box-plot represent the relative upregulation of p-STING on PBMCs in response to cfChPs treatment. **(D)** Representative fluorescent confocal microphotographs showing activation of c-GAS protein in PBMCs in response to cfChPs treatment. Internalization by PBMCs of cfChPs labelled in their histones (ATTO-488 green), triggered upregulation of c-GAS (red) protein. Quantitative analysis (MFI) reveal visible elevation of cGAS expression in the cfChPs-treated cells by 2 h and 4 h compared to untreated control cells. Results of quantitative analyses of overall cGAS activation at 2 h and 4 h and the relative presence of cGAS in the nucleus and the cytoplasm are presented as box-plots (upper and lower respectively). Results are represented as box-plot indicating median with minimum and maximum data values and the data were analyzed using two-tailed unpaired student’s *t*-tests in GraphPad Prism 8. **_*_** p<0.05, **_**_** p<0.01.

## Discussion

Several notable observations were made in this study. First, cfChPs released from dying PBMCs, as well as serum derived cfChPs, rapidly enter immune cells and accumulate within their nuclei. Second, cfChPs from both sources activate CD69 expression on T cells and NK cells; cellular stress-related pathways and inflammatory cytokines. Third, cfChPs-induced immune response is mediated via the cGAS-STING pathway.

It is currently believed that cell death results in the release of DAMPs, which are responsible for the activation of an inflammatory immune response (Denning et al., 2019; Land, 2015; Rock et al., 2011); however, the critical component(s) of DAMPs that is responsible for mounting immune response continues to remain elusive (Fleischmann et al., 2016; Schaefer, 2014; Ye et al., 2020). The composition of DAMPs is complex as they comprise a wide range of molecules derived from the extracellular matrix as well as nuclear components of dying cells (Schaefer, 2014). Our results identify that cfChPs might be a critical component of DAMPs that activate immune response following their internalization by lymphocytes. Using indirect and direct approaches, we have further demonstrated that the cfChPs-induced immune response may be mediated via the cGAS-STING pathway. For the former, we utilized a well-researched pharmacological inhibitor of STING, H151 (Al-Asmari et al., 2022; Gong et al., 2021; Hu et al., 2022, 2023; Varga et al., 2023), which prevents palmitoylation of STING and thus blocks STING signaling pathway (Haag et al., 2018). We found that H151 could abrogate cfChPs induced inflammatory cytokine release. Since the cGAS-STING pathway is activated when nucleic acids are atypically present in the cytoplasm (Motwani et al., 2019), we suspected that cfChPs activated cGAS-STING pathway while present in cytoplasm. Direct confirmation of involvement of cGAS-STING pathway came from flow cytometry and fluorescent confocal microscopy experiments. We show by flow cytometry that cfChPs treatment upregulates the phosphorylated STING protein. Experiments using fluorescent confocal microscopy demonstrated upregulation of cGAS expression in the nucleus of cfChPs treated PBMCs which appeared to shuttle back and forth between the nucleus and the cytoplasm (Herzner et al., 2021; Liu et al., 2018; Song et al., 2022). These findings suggest that cfChPs-induced immune response activation is mediated via the cGAS-STING pathway. However, further studies are required to evaluate the involvement of the cGAS-STING pathway in the cfChPs-induced immune response.

We have shown, in several studies reported elsewhere, that the various pathological activities of cfChPs can be abrogated by agents that deactivate cfChPs (Agarwal et al., 2022; Kirolikar et al., 2018; I. Mittra et al., 2017; Mittra Id et al., 2020; Ostwal et al., 2022; Pal et al., 2022; Pilankar et al., 2022). Of these, the pro-oxidant combination of the commonly used nutraceuticals resveratrol and copper (R-Cu) holds the most promise for clinical application because of its lack of toxicity and low cost. We have reported that R-Cu can ameliorate sepsis in a pre-clinical model (Mittra Id et al., 2020) and prevent cytokine storm related death in severe coronavirus disease 2019 in humans (Mittra et al., 2020). For this reason, R-Cu may be a worthy candidate for evaluation in clinical trials involving life-threatening disorders associated with hyper-inflammation, which can include sepsis, cancer, neurodegenerative disorders, and auto-inflammatory as well as autoimmune diseases.

## Materials and Methods

### Institutional ethics approval

This study was approved by the Institutional Ethics Committee (IEC) of Advanced Centre for Treatment, Research and Education in Cancer (ACTREC), Tata Memorial Centre (TMC), Mumbai (Approval no. 900520). Peripheral blood samples (10 mL) were collected from healthy volunteers after obtaining informed consent as required by the IEC.

### Collection of blood samples

Blood samples were used for isolation of PBMCs and cfChPs. To isolate PBMCs, blood samples (10 mL) were collected in heparinized Vacutainer TM tubes (Becton-Dickinson Vacutainer Systems, Franklin Lakes, NJ, USA), whereas to isolate cfChPs, samples were collected in plain Vacutainer tubes (Becton-Dickinson Vacutainer Systems). The volunteers had no history of febrile or other illnesses in the preceding 3 months.

### Isolation of PBMCs

PBMCs were isolated using Ficoll-Hypaque according to standard procedures.

### Procedure for collecting conditioned medium (CM) from hypoxia-induced dying PBMCs

ThinCert® Cell Culture Inserts, containing a separating membrane of pore size 400 nm, are designed for dual chamber experiments (Greiner Bio-One, Kremsmünster, Austria). To collect CM containing cfChPs released from dying cells, PBMCs (1 x 10^6^) were seeded in ThinCert® Cell Culture Inserts containing 1.5 mL Dulbecco’s Modified Eagle Medium (DMEM) and placed in a 35-mm 6-well plate. The plate was incubated in a hypoxia chamber (1% O_2_) for 72 h. Post 72 h, sufficient DMEM (∼700 μL) was added to the lower chamber such that the level of media touched the base of the insert placed above. The plates were incubated at 37°C in a humidified atmosphere of 5% CO_2_ for 48 h under normoxic conditions. Media in the lower chamber allowed cfChPs <400 nm in size, released from hypoxia-induced dying PBMCs, to seep into the medium in the lower chamber. Post-incubation, conditioned media from the wells was pooled and was used for further experiments.

In some experiments, PBMCs were dually fluorescently labelled in their DNA and histones such that the collected CMs contained dually fluorescently labelled cfChPs (see below).

### Collection of CM containing dually fluorescently labelled cfChPs

CM containing dually fluorescently labelled cfChPs was used in experiments to detect the uptake of fluorescently dually labelled cfChPs by recipient PBMCs. PBMCs were fluorescently dually labelled in their DNA with bromodeoxyuridine (10 μM; Sigma-Aldrich, St. Louis, MO, USA), and in their histones (H2B) with CellLight® Histone 2BGFP (Thermo Fisher Scientific, Waltham, MA, USA) for 36 h. The detailed labelling procedure has been described by us previously (Kirolikar et al., 2018). The dually labelled PBMCs were induced to undergo hypoxic cell death, as described above, such that the collected CM contained fluorescently dually labelled cfChPs (<400 nm). Two hundred micro-liters of conditioned media containing cfChPs <400nm that had seeped into the lower chamber of the insert was applied to isolated PBMCs in a time course experiment (2h, 4h and 6h). Cells were then washed and processed for fluorescence microscopy to detect presence of fluorescent signals of BrdU and histone H2BGFP in the recipient PBMCs.

### Treatment of PBMCs with CM containing cfChPs

CM collected from hypoxia-induced dying (unlabeled) PBMCs containing cfChPs were used in two experimental settings. In the first setting, a time course analysis was performed wherein PBMCs were treated with varying volumes of CMs (25, 50, 100, 200, 300, 400, and 500 µL) for estimation of CD69 expression (see below). Optimal CD69 expression was obtained using 200 µL CM. In the second experiment, PBMCs were treated with 200 µL CM to facilitate the estimation of stress markers and inflammatory cytokines (see below).

### Isolation of cfChPs from human sera

cfChPs were isolated from sera of healthy volunteers according to a protocol described by us previously (Mittra et al., 2015). Pooled serum (typically from ∼5 individuals) was used to isolate cfChPs to maintain inter-experimental consistency. Isolated cfChPs were quantified in terms of their DNA content using a Pico-Green quantification assay.

### Fluorescent dual labelling of cfChPs

Fluorescently dual-labelled cfChPs were utilized to detect the uptake of fluorescently dually labelled cfChPs by recipient PBMCs. cfChPs were dually fluorescently labelled in their DNA and histone H4 with Platinum Bright 550 (red) and ATTO-TEC 488 (green), respectively, according to the protocol described by us previously (Mittra et al., 2015).

### Treatment of PBMCs with serum derived cfChPs

PBMCs were treated with cfChPs and were analyzed for CD69 expression, cellular stress markers, and inflammatory cytokine levels. PBMCs (5 x 10^5^) were plated in 24-well plates containing 1 mL DMEM. Following overnight incubation, cells were treated with serum derived cfChPs (10 ng equivalent of DNA), with and without H151 for 24 h in a humidified atmosphere of 5% CO_2_. Untreated PBMCs were utilized as negative controls.

### qRT-PCR

qRT-PCR was used for the evaluation of stress markers, STING and cGAS after the treatment of PBMCs with 1) CM from hypoxia-induced dying PBMCs (200 µL), or 2) serum derived cfChPs (10 ng). The time points for both analyses were: 0, 6, 12, 24, 36, 48, and 72 h and appropriate untreated control PBMCs were analyzed in parallel. Total RNA was isolated using RNeasy Mini Kit (Qiagen, Hilden, Germany), and approximately 1 µg of isolated RNA was converted to cDNA using a RT^2^ First Strand Kit (Qiagen, Hilden, Germany). qRT-PCR was performed using a SYBR Select Master Mix (Applied Biosystems, Waltham, MA, USA). All samples were assayed on a QuantStudio5 system (Applied Biosystems) in duplicates. Data were analyzed using a comparative C_T_ method, and fold change in mRNA expression was calculated as 2^(-ΔΔC_T_).

### Flow cytometry

Flow cytometry was utilized for the evaluation of 1) CD69 expression, 2) inflammatory cytokine levels, and 3) stress-related markers and p-STING following the treatment of PBMCs with either CM (200 µL) or serum derived cfChPs (10 ng).

#### CD69 expression

After 24 h incubation with CM or cfChPs, PBMCs were labelled for 20 min with anti-CD45-BUV395 (HI30), anti-CD3-BV711 (UCHT1), anti-CD56-PE-Cy7 (NCAM) and anti-CD69-APC (FN50) (Becton Dickinson Biosciences, San Jose, CA, USA.) in dark at room temperature. Samples were analyzed on a FACSAria III cytometer (Becton Dickinson).

#### Estimation of cytokine levels

After 48 h treatment either with CM or with cfChPs, cultured cells were centrifuged at 3,000 rpm for 10 min at 4^°^C and supernatants were collected and stored at -80°C for cytokine analysis. A LEGENDplex™ Multi-Analyte Flow assay kit (BioLegend, San Diego, CA, USA) or ELISA was used as per the manufacturer’s instructions to quantify levels of secreted cytokines in culture supernatants.

#### Stress-related markers and pSTING expression

After incubation with cfChPs at the peak time-point as detected by qRT-PCR, PBMCs were fixed and permeabilized with either 4% formaldehyde and 90% methanol or cytofix/cytoperm kit (Becton Dickinson) based on the manufacturer’s instructions and were then stained using specific primary antibodies against respective proteins. For unconjugated antibodies, cells were stained with appropriate secondary antibody. Cells were acquired on Attune NxT flow cytometer (ThermoFisher Scientific, USA).

#### Flow cytometry acquisition

To determine CD69 expression, lymphocytes were gated on forward and side scatter parameters and further as CD45^+^ cells **(Suppl. Fig. 5).** T-cells were gated as CD3^+^ cells and NK-cells as CD56^+^CD3^-^ cells. CD69 expression on the cells was determined as percentage of positive cells in the respective gated cell regions. The threshold between negative and positive was defined by the fluorescence minus one method. At least 10,000 lymphocytes were acquired on a FACSAria III cytometer (Becton Dickinson). To determine stress markers and pSTING expression, lymphocytes were gated on forward and side scatter parameters and specific marker expression on the cells was determined by increase in Mean fluorescence intensity (MFI) compared to only secondary antibody staining or only cells (whichever applicable). At least 5,000 lymphocytes were acquired on Attune NxT flow cytometer (ThermoFisher Scientific, USA). For the estimation of cytokine levels, sample acquisition was performed as per the manufacturer’s instructions. At least 4,000 cytometric beads were acquired on a FACSAria III cytometer (Becton Dickinson).

#### Flow cytometry analysis

Data was analyzed using FlowJo software (Version 10; FlowJo LLC, Ashland, OR, USA) and cytokine data was analyzed using LEGENDplex Data Analysis Software (BioLegend).

### Enzyme-linked immunosorbent assay (ELISA)

ELISA was used to estimate IL-1β, IFN-β, and CXCL9 levels in the supernatant media (Krishgen Biosystems, Mumbai, India) as per the manufacturer’s instructions.

### Fluorescent confocal microscopy

For co-staining of cfChPs and cGAS protein, cfChPs were labelled only in their histones with ATTO-488 (green) to prevent spectral overlap with cGAS staining. PBMCs were treated with labelled cfChPs and incubated for 2 h and 4 h at 37°C in a humidified atmosphere of 5% CO_2_. Cells were cytospun onto a slide, immediately fixed with 4% PFA and immuno-stained using rabbit anti-cGAS primary antibody and appropriate secondary antibody by indirect immunofluorescence method as described by us earlier (Indraneel Mittra et al., 2017). Slides were mounted in VectaShield DAPI (Vector labs, USA). For confocal microscopy, images were acquired using a 63x oil objective on Nikon AX Confocal Microscope System, (Nikon Corporation, Japan). All experiments were performed in duplicate, and 50 cells were analysed in each case. Mean fluorescence intensity (MFI) of images was measured using ImageJ FIJI software 1.5t version (W. Rasband, Bethesda, USA) and results were expressed as mean ± SEM.

### Statistical analysis

Statistical analysis was performed using two-tailed unpaired student’s *t*-tests using GraphPad Prism 8 (GraphPad Software, Boston, MA, USA) and the results are expressed as mean ± SEM. The following significance thresholds were applied: **_*_** p<0.05, **_**_** p<0.01, _***_ p<0.005, and _****_ p<0.0001.

## Data Availability

All data supporting the findings of this study are available within the article and supplementary materials.

## Acknowledgments

The authors thank personnel in the flow cytometry facility at ACTREC-TMC for their technical support. The authors thank Dr. Rajan Basak for his guidance in qRT-PCR.

## Supplementary Files

**Suppl. Fig. 1:** Accumulation of cfChPs released from dying PBMCs in the nuclei of healthy PBMCs. PBMCs were dually labelled in their DNA and histone with bromodeoxyuridine (red) and Histone2B-GFP (green), respectively, and induced to undergo hypoxic cell death. The filtered CM (200 µL) containing dually labelled cfChPs (<400 nm) were applied to fresh PBMCs as described in the Materials and Methods section. The upper panel represents the images of dually labelled donor PBMCs; the lower panel represents treated recipient PBMCs. Fluorescent microscopy images were taken at 4 h.

**Suppl. Fig. 2:** Upregulation of stress-related markers in human PBMCs following treatment with CM (200μl). mRNA expression of six stress-related markers in PBMCs was detected by qRT-PCR and analyzed using a comparative C_T_ method. Line graphs represent the results of time course analyses calculated as expression fold [2^(-ΔΔC_T_)].

**Suppl. Fig. 3:** Accumulation of cfChPs in the nuclei of PBMCs. Serum derived cfChPs were dually labelled in their DNA and histones with Platinum Bright 550 (red) and ATTO-TEC-488 (green), respectively. PBMCs were treated with the dually labelled cfChPs (10 ng) and fluorescent microscopy images were taken at 2 h.

**Suppl. Fig. 4:** Upregulation of stress-related markers in human PBMCs following treatment with cfChPs (10 ng). mRNA expression of six stress-related markers in PBMCs was detected by qRT-PCR and analyzed using a comparative C_T_ method. Line graphs represent the results of time course analyses calculated as expression fold [2^(-ΔΔC_T_)].

**Suppl. Fig. 5:** Flow cytometry plots representing gating strategy. Gating of lymphocytes on forward/side scatter and further as CD45+ cells, T-cells were identified as CD3+ cells and NK cells as CD56+/CD3- cells.

## References

Agarwal A, Khandelwal A, Pal K, Khare NK, Jadhav V, Gurjar M, Punatar S, Gokarn A, Bonda A, Nayak L, Kannan S, Gota V, Khattry N, Mittra I. 2022. A novel pro-oxidant combination of resveratrol and copper reduces transplant related toxicities in patients receiving high dose melphalan for multiple myeloma (RESCU 001). PLoS One 17:e0262212. doi:10.1371/journal.pone.0262212

Akira S, Uematsu S, Takeuchi O. 2006. Pathogen recognition and innate immunity. Cell. doi:10.1016/j.cell.2006.02.015

Al-Asmari SS, Rajapakse A, Ullah TR, Pépin G, Croft L V., Gantier MP. 2022. Pharmacological Targeting of STING-Dependent IL-6 Production in Cancer Cells. Front Cell Dev Biol 9. doi:10.3389/fcell.2021.709618

Ashida H, Mimuro H, Ogawa M, Kobayashi T, Sanada T, Kim M, Sasakawa C. 2011. Cell death and infection: A double-edged sword for host and pathogen survival. J Cell Biol. doi:10.1083/jcb.201108081

Chaplin DD. 2010. Overview of the immune response. J Allergy Clin Immunol 125:S3. doi:10.1016/j.jaci.2009.12.980

Denning NL, Aziz M, Gurien SD, Wang P. 2019. Damps and nets in sepsis. Front Immunol. doi:10.3389/fimmu.2019.02536

Fleischmann C, Scherag A, Adhikari NKJ, Hartog CS, Tsaganos T, Schlattmann P, Angus DC, Reinhart K. 2016. Assessment of global incidence and mortality of hospital-treated sepsis current estimates and limitations. Am J Respir Crit Care Med 193:259–272. doi:10.1164/rccm.201504-0781OC

Gong W, Lu L, Zhou Y, Liu J, Ma H, Fu L, Huang S, Zhang Y, Zhang A, Jia Z. 2021. The novel STING antagonist H151 ameliorates cisplatin-induced acute kidney injury and mitochondrial dysfunction. Am J Physiol Physiol 320:F608– F616. doi:10.1152/ajprenal.00554.2020

Haag SM, Gulen MF, Reymond L, Gibelin A, Abrami L, Decout A, Heymann M, van der Goot FG, Turcatti G, Behrendt R, Ablasser A. 2018. Targeting STING with covalent small-molecule inhibitors. Nature 559:269–273. doi:10.1038/s41586-018-0287-8

Herzner A-M, Schlee M, Bartok E. 2021. The many faces of cGAS: how cGAS activation is controlled in the cytosol, the nucleus, and during mitosis. Signal Transduct Target Ther 6:260. doi:10.1038/s41392-021-00684-3

Hu S, Gao Y, Gao R, Wang Y, Qu Y, Yang J, Wei X, Zhang F, Ge J. 2022. The selective STING inhibitor H-151 preserves myocardial function and ameliorates cardiac fibrosis in murine myocardial infarction. Int Immunopharmacol 107:108658. doi:10.1016/j.intimp.2022.108658

Hu Z, Zhang F, Brenner M, Jacob A, Wang P. 2023. The protective effect of H151, a novel STING inhibitor, in renal ischemia-reperfusion-induced acute kidney injury. Am J Physiol Physiol 324:F558–F567. doi:10.1152/ajprenal.00004.2023

Kirolikar S, Prasannan P, Raghuram G V., Pancholi N, Saha T, Tidke P, Chaudhari P, Shaikh A, Rane B, Pandey R, Wani H, Khare NK, Siddiqui S, D’souza J, Prasad R, Shinde S, Parab S, Nair NK, Pal K, Mittra I. 2018. Prevention of radiation-induced bystander effects by agents that inactivate cell-free chromatin released from irradiated dying cells. Cell Death Dis 9:1–16. doi:10.1038/s41419-018-1181-x

Land WG. 2015. The role of damage-associated molecular patterns in human diseases: Part I - Promoting inflammation and immunity. Sultan Qaboos Univ Med J.

Liu H, Zhang H, Wu X, Ma D, Wu J, Wang L, Jiang Y, Fei Y, Zhu C, Tan R, Jungblut P, Pei G, Dorhoi A, Yan Q, Zhang F, Zheng R, Liu S, Liang H, Liu Z, Yang H, Chen J, Wang P, Tang T, Peng W, Hu Z, Xu Z, Huang X, Wang J, Li H, Zhou Y, Liu F, Yan D, Kaufmann SHE, Chen C, Mao Z, Ge B. 2018. Nuclear cGAS suppresses DNA repair and promotes tumorigenesis. Nature 563:131–136. doi:10.1038/s41586-018-0629-6

Mittra I, de Souza R, Bhadade R, Madke T, Shankpal PD, Joshi M, Qayyumi B, Bhattacharya A, Gota V, Gupta S, Chaturvedi P, Badwe R. 2020. Resveratrol and Copper for treatment of severe COVID-19: an observational study (RESCU 002) (preprint). medRxiv 2020.07.21.20151423.

Mittra I, Khare NK, Raghuram GV, Chaubal R, Khambatti F, Gupta D, Gaikwad A, Prasannan P, Singh Akshita, Iyer A, Singh Ankita, Upadhyay P, Nair NK, Mishra PK, Dutt A. 2015. Circulating nucleic acids damage DNA of healthy cells by integrating into their genomes. J Biosci 40:91–111. doi:10.1007/s12038-015-9508-6

Mittra I., Pal K, Pancholi N, Shaikh A, Rane B, Tidke P, Kirolikar S, Khare NK, Agrawal K, Nagare H, Nair NK. 2017. Prevention of chemotherapy toxicity by agents that neutralize or degrade cell-free chromatin. Ann Oncol 28:2119– 2127. doi:10.1093/annonc/mdx318

Mittra Indraneel, Samant U, Sharma S, Raghuram G V, Saha T, Tidke P, Pancholi N, Gupta D, Prasannan P, Gaikwad A, Gardi N, Chaubal R, Upadhyay P, Pal K, Rane B, Shaikh A, Salunkhe S, Dutt S, Mishra PK, Khare NK, Nair NK, Dutt A. 2017. Cell-free chromatin from dying cancer cells integrate into genomes of bystander healthy cells to induce DNA damage and inflammation. Cell Death Discov 3:17015. doi:10.1038/cddiscovery.2017.15

Mittra Id I, Pal K, Pancholi N, Tidke P, Siddiqui S, Rane B, D’souza J, Shaikh A, Parab S, Shinde S, Jadhav V, Shende S, Raghuram G V. 2020. Cell-free chromatin particles released from dying host cells are global instigators of endotoxin sepsis in mice. PLoS One 15(3):e0229017. doi:10.1371/journal.pone.0229017

Motwani M, Pesiridis S, Fitzgerald KA. 2019. DNA sensing by the cGAS–STING pathway in health and disease. Nat Rev Genet 20:657–674. doi:10.1038/s41576-019-0151-1

Ostwal V, Ramaswamy A, Bhargava P, Srinivas S, Mandavkar S, Chaugule D, Peelay Z, Baheti A, Tandel H, Jadhav VK, Shinde S, Jadhav S, Gota V, Mittra I. 2022. A pro-oxidant combination of resveratrol and copper reduces chemotherapy-related non-haematological toxicities in advanced gastric cancer: results of a prospective open label phase II single-arm study (RESCU III study). Med Oncol 40:17. doi:10.1007/s12032-022-01862-1

Pal K, Raghuram G V., Dsouza J, Shinde S, Jadhav V, Shaikh A, Rane B, Tandel H, Kondhalkar D, Chaudhary S, Mittra I. 2022. A pro-oxidant combination of resveratrol and copper down-regulates multiple biological hallmarks of ageing and neurodegeneration in mice. Sci Rep 12:17209. doi:10.1038/s41598-022-21388-w

Pilankar A, Singhavi H, Raghuram G V., Siddiqui S, Khare NK, Jadhav V, Tandel H, Pal K, Bhattacharjee A, Chaturvedi P, Mittra I. 2022. A pro-oxidant combination of resveratrol and copper down-regulates hallmarks of cancer and immune checkpoints in patients with advanced oral cancer: Results of an exploratory study (RESCU 004). Front Oncol 12. doi:10.3389/fonc.2022.1000957

Rock KL, Lai JJ, Kono H. 2011. Innate and adaptive immune responses to cell death. Immunol Rev 243:191–205. doi:10.1111/j.1600-065X.2011.01040.x

Schaefer L. 2014. Complexity of danger: The diverse nature of damage-associated molecular patterns. J Biol Chem. doi:10.1074/jbc.R114.619304

Shabrish S, Mittra I. 2021. Cytokine Storm as a Cellular Response to dsDNA Breaks: A New Proposal. Front Immunol 12. doi:10.3389/fimmu.2021.622738

Song J-X, Villagomes D, Zhao H, Zhu M. 2022. cGAS in nucleus: The link between immune response and DNA damage repair. Front Immunol 13. doi:10.3389/fimmu.2022.1076784

Varga KZ, Gyurina K, Radványi Á, Pál T, Sasi-Szabó L, Yu H, Felszeghy E, Szabó T, Röszer T. 2023. Stimulator of Interferon Genes (STING) Triggers Adipocyte Autophagy. Cells 12:2345. doi:10.3390/cells12192345

Wang H, Shan X, Ren M, Shang M, Zhou C. 2021. Nucleosomes enter cells by clathrin- and caveolin-dependent endocytosis. Nucleic Acids Res 49:12306–12319. doi:10.1093/nar/gkab1121

Ye Q, Wang B, Mao J. 2020. The pathogenesis and treatment of the ‘Cytokine Storm’’ in COVID-19.’ J Infect. doi:10.1016/j.jinf.2020.03.037

