## Supplementary figures and images for "Cell-free chromatin particles from dying cells promote the induction of an immune response in human lymphocytes"

### Supplementary Figure 1

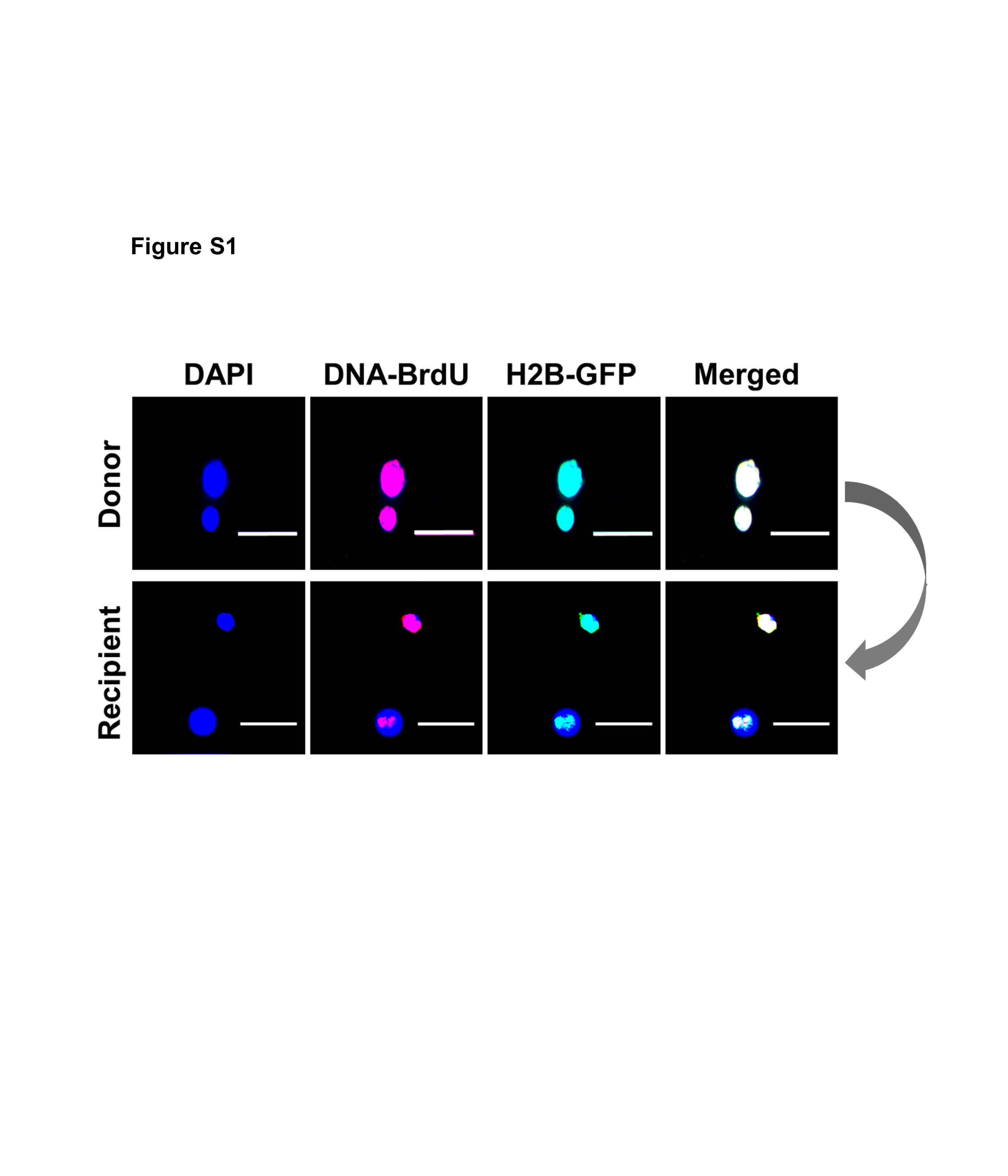

### Supplementary Figure 2

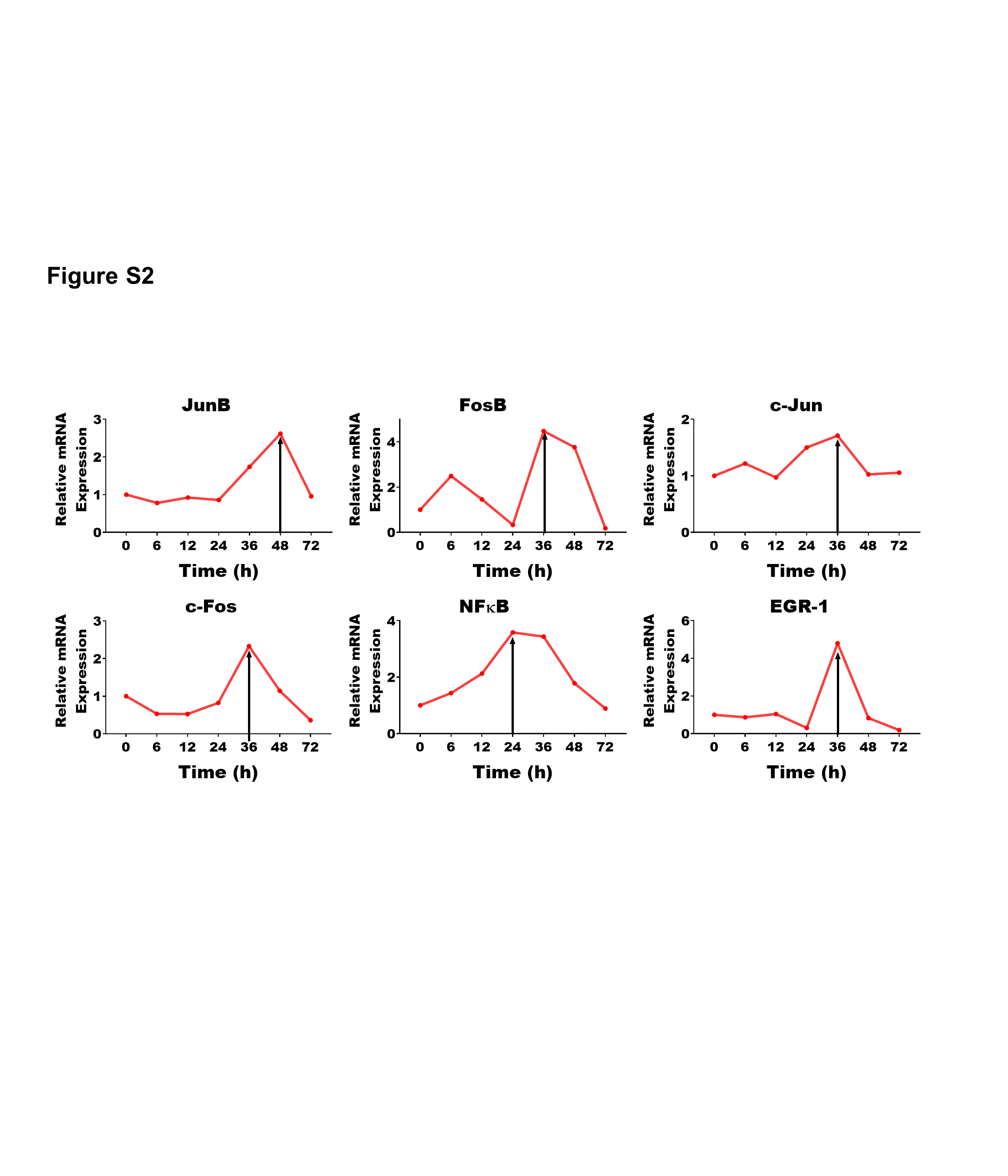

### Supplementary Figure 3

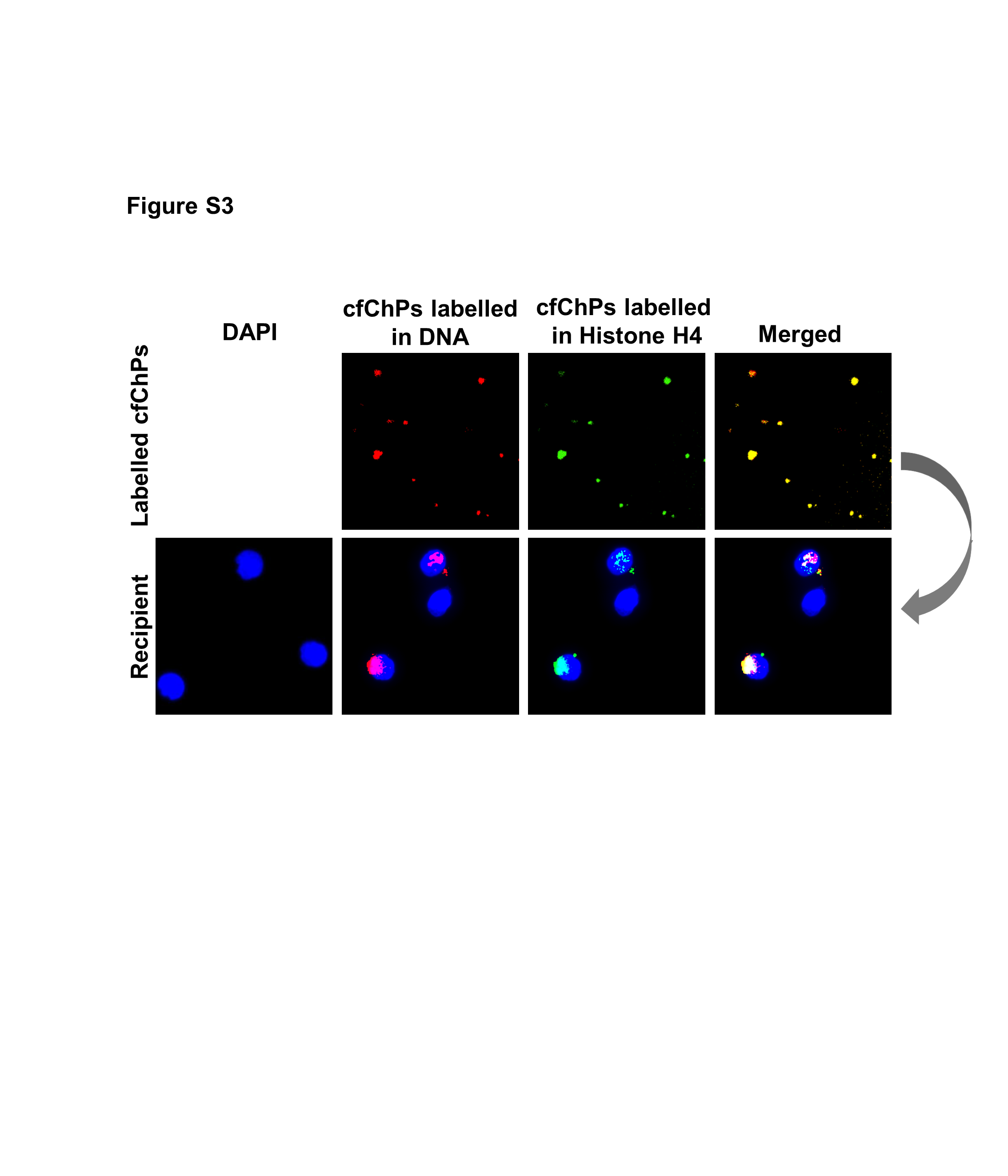

### Supplementary Figure 4

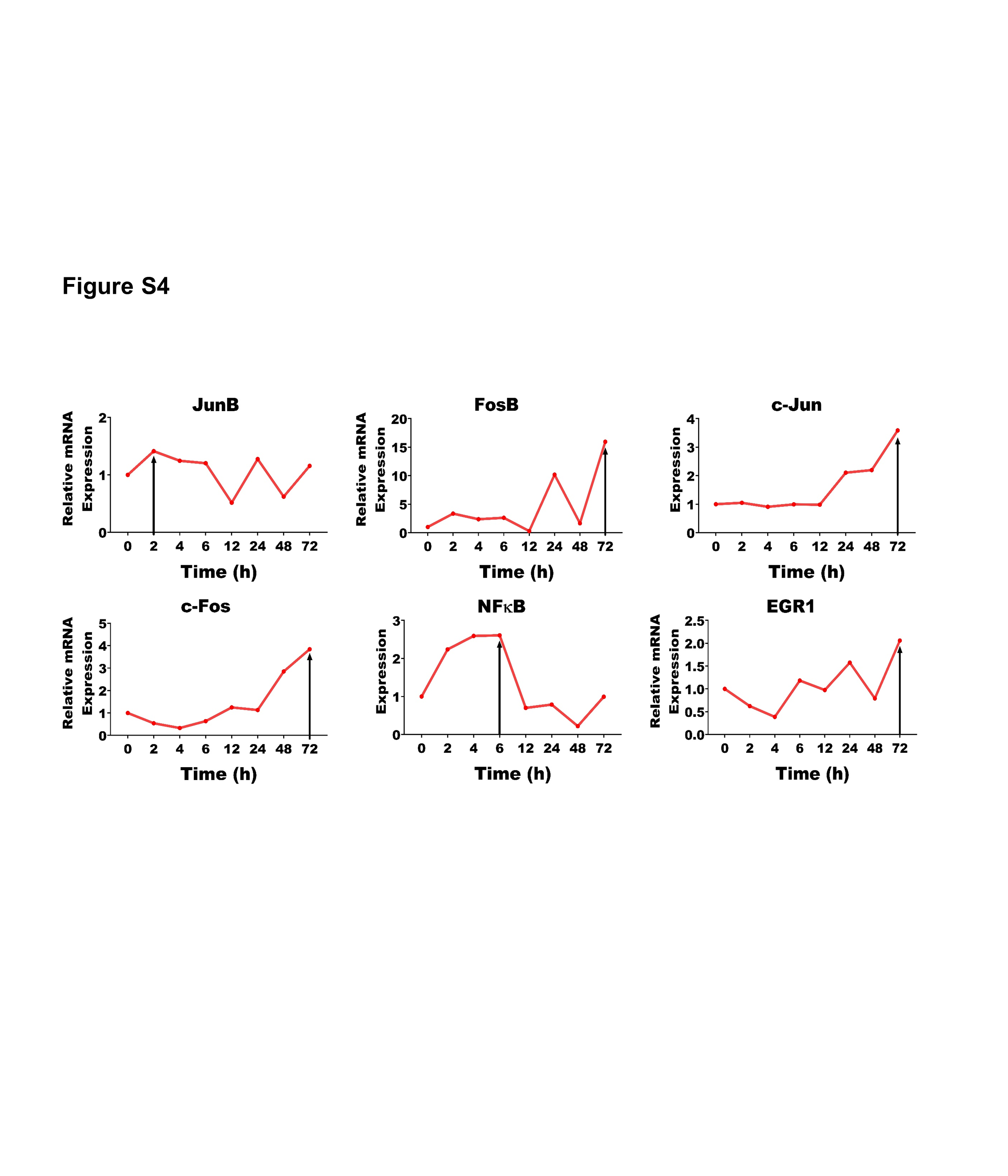

### Supplementary Figure 5

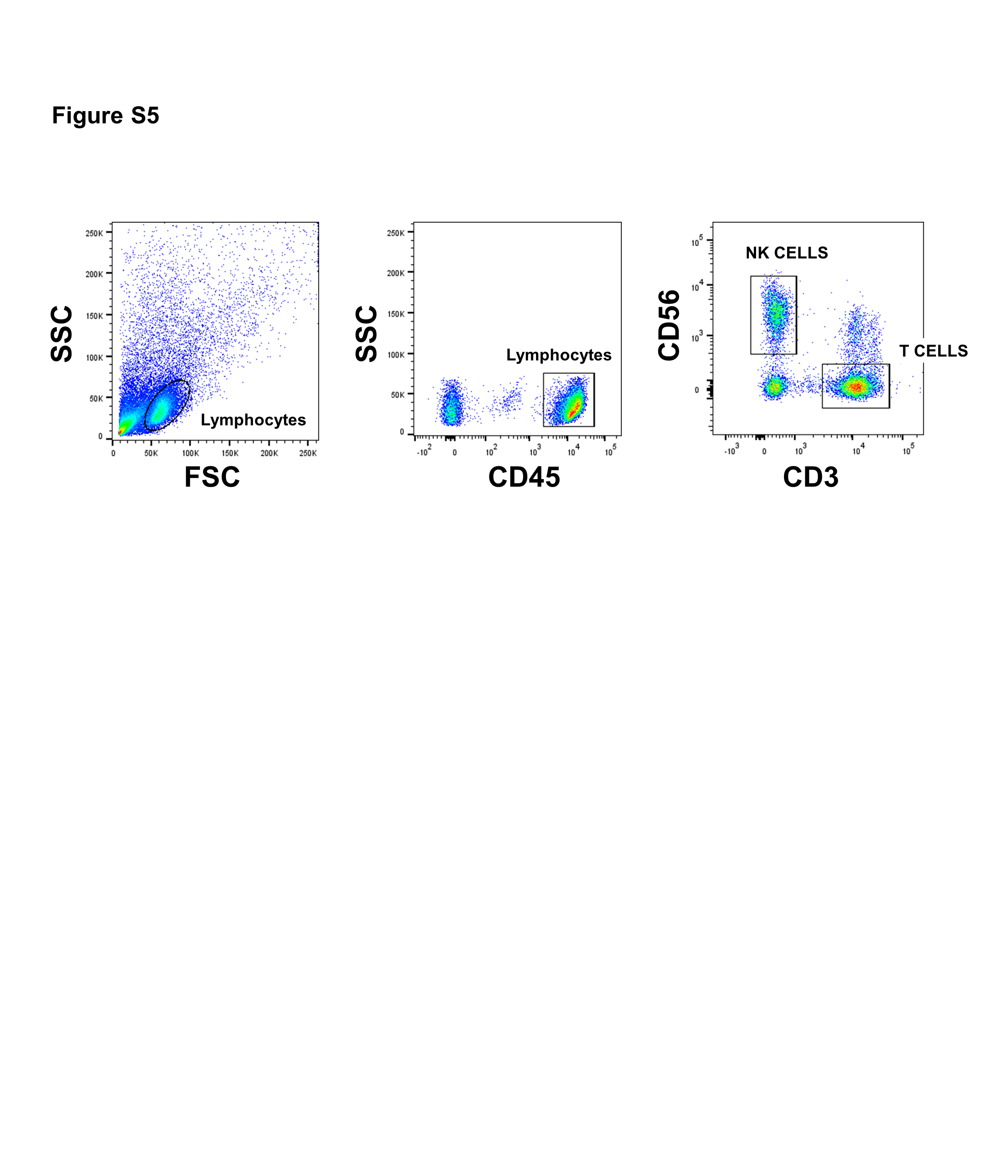
